# Genomic perspectives on adaptation and conservation in the endangered long-tailed goral (*Naemorhedus caudatus*)

**DOI:** 10.1101/505156

**Authors:** Jessica A. Weber, Oksung Chung, Jungeun Kim, JeHoon Jun, Hyejin Lee, Sungwoong Jho, Yun Sung Cho, Dae-Soo Kim, Woon ki Paek, Soonok Kim, Hanshin Lee, Semin Lee, Jeremy S. Edwards, Joseph A. Cook, Junghwa An, Jong Bhak

**Affiliations:** Department of Biology and Museum of Southwestern Biology, University of New Mexico, Albuquerque, NM 87131, United States; Personal Genomics Institute, Genome Research Foundation, Osong 28160, Republic of Korea; Geromics, Ulsan 44919, Republic of Korea; The Genomics Institute (TGI), Ulsan National Institute of Science and Technology (UNIST), Ulsan 44919, Republic of Korea; Rare Disease Research Center, Korea Research Institute of Bioscience and Biotechnology, Daejeon, 305-333, Republic of Korea; National Science Museum, Daejeon, 34143, Republic of Korea; National Institute of Biological Resources, Incheon, 22689, Republic of Korea; University of California - San Diego, CA 92122, United States; Department of Biomedical Engineering, School of Life Sciences, UNIST, Ulsan, 44919, Republic of Korea; Department Chemistry and Chemical Biology, Molecular Genetics and Microbiology, and Chemical and Nuclear Engineering, University of New Mexico - Albuquerque, NM 87131, United States

**Keywords:** Caprini, heterozygosity, historical demography, positively selected genes

## Abstract

Native to the mountains of East Asia, the long-tailed goral (*Naemorhedus caudatus*) is a vulnerable wild ungulate in the tribe Caprini. To understand key conservation issues related to fragmentation and subsequent endangerment of this montane species, we sequenced and analyzed the genome of a long-tailed goral to explore historical demography, contemporary levels of genetic diversity, and potential immune response. When compared to ten additional mammalian reference genomes, we identified 357 positively selected genes (PSGs) in the long-tailed goral and 364 PSGs in the Caprini lineage. Gene Ontology analyses showed statistical enrichment in biological processes related to immune function and in genes and pathways related to blood coagulation. We also identified low levels of heterozygosity (0.00114) in the long-tailed goral along with decreases in population size relative to other species of Caprini. Finally, we provide evidence for positive selection on the muscle development gene *myostatin* (*MSTN*) in the Caprini lineage, which may have increased the muscle development and climbing ability in the caprine common ancestor. Low effective population size and decreased heterozygosity of the long-tailed goral raise conservation concern about the effects of habitat fragmentation, over harvest, and inbreeding on this vulnerable species.

## Introduction

The long-tailed goral (*Naemorhedus caudatus*), also known as the Amur goral, is a small goat-like ungulate that has been classified as Vulnerable by the International Union for Conservation of Nature (IUCN) and Endangered by the South Korean Ministry of Environment ^1^. While bovid taxonomy has been historically challenging to resolve, gorals are now recognized as members of the Caprini tribe of the subfamily Antilopinae, within the diverse and widespread Bovidae family. Caprines are thought to have originated in the late Miocene roughly 11 million years ago (MYA) in the mountainous archipelago between the Mediterranean and Paratethys Seas ^2^. About that time, the caprine common ancestor evolved shortened metacarpals ideally suited for climbing, allowing it to colonize new ecological niches associated with steep mountains that were not occupied by other large herbivores ^2^. As a result, caprines rapidly radiated across the mountain ranges of the Northern Hemisphere and now include 62 species in 14 genera ^3^.

The genus *Naemorhaedus* is comprised of six species of gorals that inhabit the mountains of Asia ^3,4^. Long-tailed gorals are adept climbers that inhabit steep rocky slopes ^1^ and are found in the rugged mountainous forests and shrublands of the Korean peninsula, eastern Russia (Primorsky and Khabarovsk Territories), and northeastern China ^5^. In Korea, the long-tailed goral ranges throughout the Taebaek and Sobaek mountains, spanning elevations from sea level to 2,000 m. They are diurnal browsers that feed on grasses, shoots, leaves, nuts, and fruits and occur in small groups of 4-12 individuals with a group home range of roughly 40 hectares ^1^. Previous studies of goral populations in Korea have shown a preference for steep slopes as a defense against predators, though they were observed to descend to lower habitats during periods of heavy winter snow when vegetation is unavailable at higher elevations ^6-8^. Various serious diseases have been described in both wild and captive populations of long-tailed gorals ^9^, and deaths due to taeniasis parasites (tapeworms), pneumonia, gastroenteritis, and hepatitis have been reported in captive individuals ^10^. Surveys of a Russian population of long-tailed gorals identified more than 30 species of helminth parasites ^11-13^, which result in massive infections that can cause the death of the host, as occurred in 1,937 when all ten long-tailed gorals sent from the Lazovsky Nature Reserve to the Moscow Zoo died within eight months of their arrival ^11^. In addition, extremely high gastrointestinal helminth parasite loads were also reported in closely related *N. goral* captive individuals, and mixed endoparasite infections were seen in the majority of the wild animals examined ^14^.

The long-tailed goral was internationally classified as Vulnerable in 2008 due to over-exploitation by hunters and habitat destruction and degradation ^1,15^, and is protected in the Republic of Korea as both an endangered wild species and as a designated national monument through the Cultural Heritage Administration ^15^. Despite its protected status, the long-tailed goral continues to be poached for meat and traditional medicine ^1^. Its range has significantly decreased over the last century, with the majority of individuals now occupying the Russian and Korean coast of the Sea of Japan ^4^. Deforestation from timber operations, along with increased human encroachment and agricultural expansion, have fragmented populations and heightened concerns of inbreeding depression and reduced effective population size ^16-18^. Population surveys in 2002 found a significant decline in the South Korean population with a total of 690-784 extant individuals ^19^. Microsatellite studies of South Korean long-tailed goral populations have shown populations along the Taebaek Mountains to be genetically distinct with lower genetic diversity in wild individuals from the southern group relative to the northern group ^20^. Furthermore, Russian surveys in the early 20^th^ century estimated roughly 2,000 individuals, which have since been reduced to about 900 individuals ^4,21^. The species is believed to be almost extinct in China ^22,23^. Across its range, it is now missing from many areas of its original occurrence; and only six subpopulations currently have more than 100 individuals, while many of the other localities only contain a few animals ^4^. Taken together, the recent range loss and population declines raise heightened concerns about the future of this species.

To better understand genome evolution and the demographic history of the long-tailed goral, and to provide a foundation for future conservation studies, we generated and analyzed the whole genome sequence (WGS) of a wild individual from the northern Taebaek Mountains in the Republic of Korea. We compared the genetic diversity of the long-tailed goral to other close species of Caprini and then performed demographic modeling to infer historical population structure. We also compared the long-tailed goral genome to the reference genomes of seven other bovids (goat, sheep, mouflon, Tibetan antelope, bison, cattle, and yak) and three distantly related mammals (human, pig, and horse) to characterize environmental adaptations present in the long-tailed goral and the caprine common ancestor.

## Results and Discussion

### Long-tailed goral whole genome sequencing

A blood sample of a wild long-tailed goral from the northern range of the Taebaek Mountains in the Republic of Korea (Fig. 1) was collected through the Association of Korean Goral Conservation. Genomic DNA was sequenced using the Illumina HiSeq2000 platform, from which we obtained 430 million (M) paired reads (Additional file 1: Table S1). The reads were then mapped to the closely related sheep (*Ovis aries*) reference genome. In total, 94% of the WGS reads were aligned to the sheep genome and yielded a 27x depth of coverage. Using the aligned reads, a variant analysis identified 55 M single nucleotide variants (SNVs) between the long-tailed goral and the sheep. To quantify the genomic effects of possible inbreeding depression resulting from habitat fragmentation and isolation in this species, we examined the proportion of heterozygous SNVs. In total, the variant composition included 52,621,471 (94.71%) homozygous and 2,936,977 (5.29%) heterozygous SNVs. We also analyzed the number of homozygous and heterozygous SNVs for each chromosome (Additional file 1: Table S2). The average homozygous and heterozygous SNVs for each chromosome were 21.12/Kb (20.12-22.34) and 1.29/Kb (0.93-1.86), respectively. Higher number of SNVs was observed in autosomes than in the sex chromosome (Chr. X). We also analyzed the homozygous SNV hotspots with 0.5M windows. Hot spots were defined by Z-score > 2 (Additional file 2: Figure S1, pink) and Z-score > 3 (red). Most of the hotspots were distributed on the chromosome ends. The heterozygous SNVs were replaced by “N” and the homozygous SNVs were used to construct a 2.61 Gb consensus sequence, which we used to predict 18,690 genes with 98.6% of the coding sequence (CDS) regions having a read depth greater than or equal to five.

**Figure 1.**
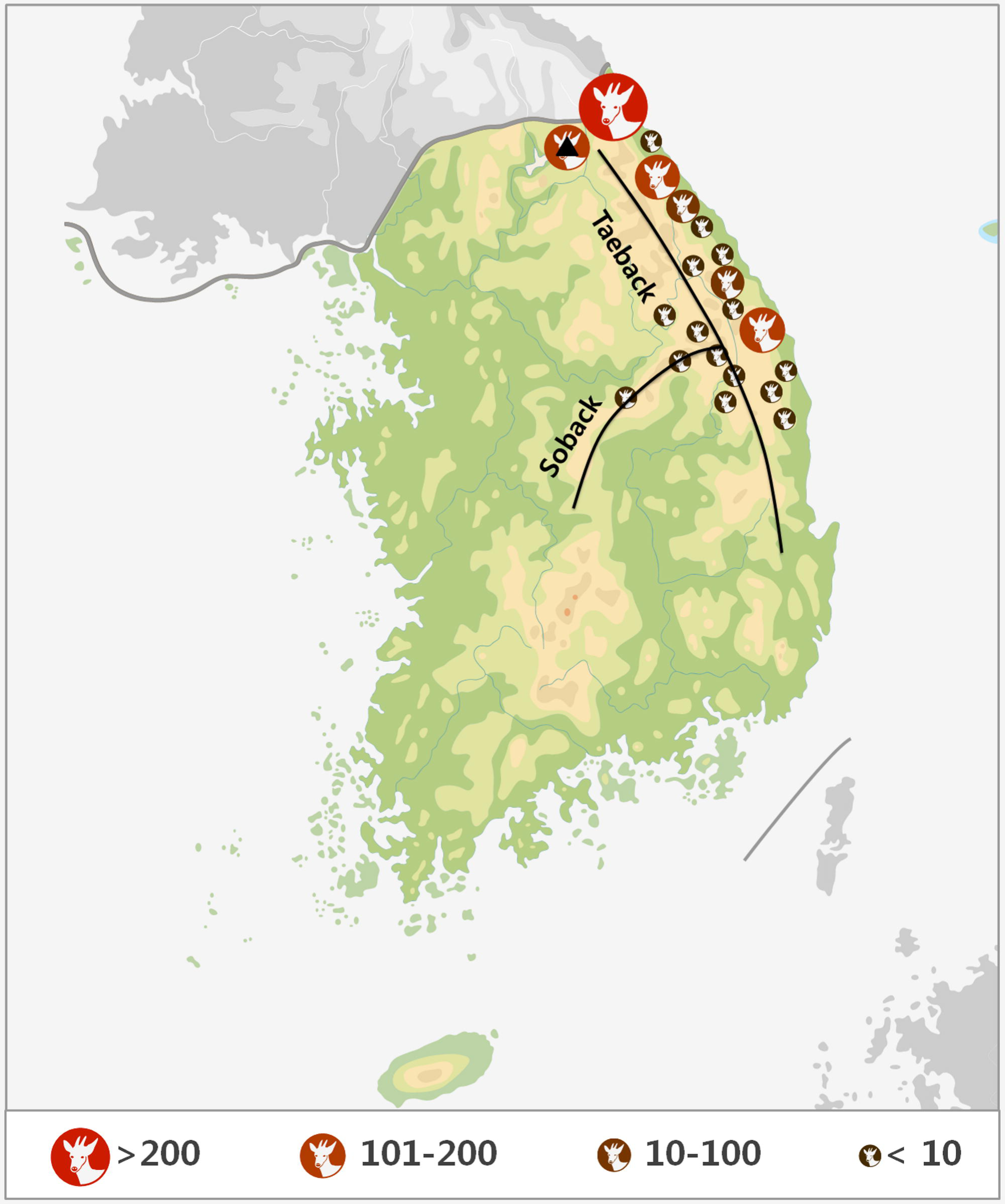
Distribution of the long-tailed goral across the Korea peninsula. The circles on the map represent population densities of the long-tailed goral which were surveyed in 2002 by the Ministry of the Environment, Korea. The black lines represent the Taback and Soback mountains which are representative habitats of the long-tailed goral. The black triangle is the location of the Association of Korea Goral Conservation, where we sampled.

### Evolutionary relationships of the long-tailed goral inferred by phylogenomic analysis

For comparative evolutionary analyses, ten mammalian reference genomes were downloaded and analyzed to infer the evolutionary history of the long-tailed goral. Included were four additional members of the Caprini tribe in the Antilopinae subfamily (goat, sheep, mouflon, and Tibetan antelope), three members of another bovid subfamily, Bovinae, in the tribe Bovini (bison, cow, and yak), and three distantly related mammals (horse, pig, and human). To reveal evolutionary relationships among the orthologous genes, we constructed orthologous gene clusters with these ten reference mammalian genomes. A total of 28,661 ortholog clusters were identified, with 7,533 (26.3%) clusters shared by all ten of the reference mammals. Of these, 5,279 (70.1%) were also single-copy long-tailed goral orthologs. Divergence times were calculated using 656,303 fourfold degenerate sites in the single-copy orthologous gene families (Fig. 2). Phylogenomic analyses showed a deep divergence of the Bovidae from the Suidae roughly 57.9 million years ago (MYA), consistent with previous findings ^24^. Divergence of the Antilopinae and Bovinae subfamilies within Bovidae was dated to 14.1 MYA and is close to previous estimates (17.6-15.1 MYA) ^25^. The Tibetan antelope is the sole species in the genus *Pantholops* and its classification has historically been disputed ^26^. Recent genetic work has reassigned it from Antilopini, where it was traditionally placed, to the Caprini tribe where it is sister to all other caprine species ^3,5,27,28^. While our phylogeny did not include genomes from other Antilopinae tribes, we estimated that the Caprini (goat, sheep, mouflon, and Tibetan antelope) shared a most recent common ancestor (MRCA) 6.38 MYA. That date is notable because it is much younger than the proposed 14 MYA split between the Caprini and Antilopini ^25^, lending genomic support to the previous genetic-based assignment of *Pantholops* to Caprini.

### Positively selected genes in the long-tailed goral and caprines

To gain insight into potential environmental adaptations that evolved in the long-tailed goral and in the Caprini common ancestor, we analyzed the 5,279 single-copy orthologous genes across our eleven species to infer genes that experienced positive selection in these lineages. In the long-tailed goral, we identified 218 and 153 PSGs by applying the branch and branch-site models, respectively (Additional file 1: Table S3 and S4). Gene Ontology (GO) enrichment analyses of the long-tailed goral PSGs against the Caprini species *Ovis aires* (sheep) as a background revealed nine biological processes related to immune function (Additional file 1: Table S5 and S6). In addition, six KEGG pathways related to immune function were found to be statistically enriched in the long-tailed goral (Additional file 1: Table S7 and S8). Previous analyses of evolutionary rate differences across mammalian genomes have reported rapid evolution in immune function, particularly in genes related to the innate immune response ^29^.

**Figure 2.**
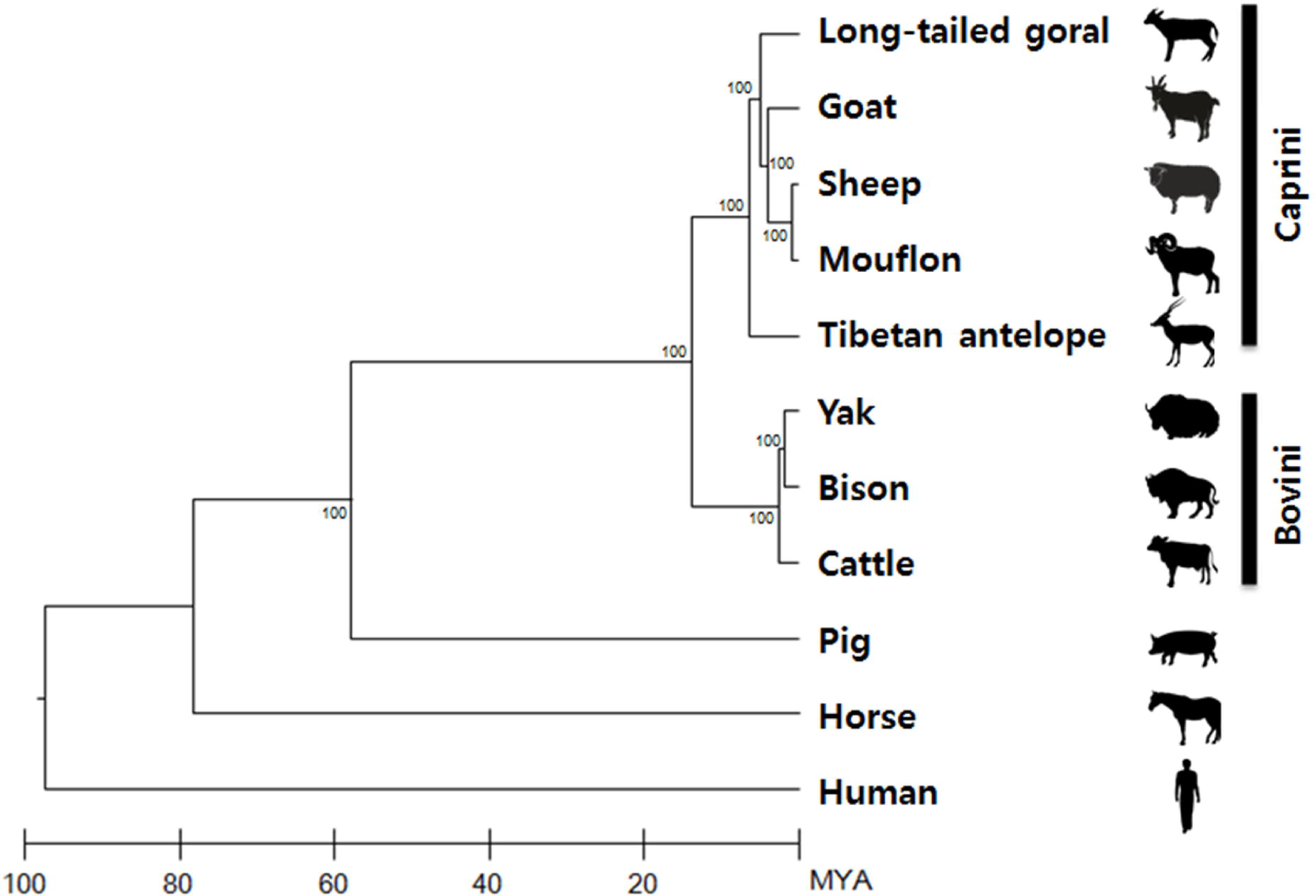
Phylogenetic relationships and divergence times. The phylogenetic topology was constructed with the 5,279 single copy orthologues defined in this study using RAxML (bootstrap support listed at nodes). The divergence times were estimated using four-fold degenerate sites of one-to-one ortholog families using the program MEGA7.

A critical component of the immune response, Type 2 T helper cells (T_H_2) are believed to have evolved to combat extracellular parasites and are the primary immune response to helminthiasis ^30-33^, which has been shown to cause mortality among captive gorals ^14^. Among the T_H_2 PSGs identified in the long-tailed goral, *interleukin 5* (*IL5*) is a T_H_2 cytokine known to play critical roles in the immune response to extracellular parasitic infections. Numerous studies of IL5 mouse knockouts have shown this cytokine to be critical to both primary and secondary immune responses to helminthiasis from a variety of nematode species ^34-36^. Also of interest, the PSG *C-C motif chemokine ligand 25* (*CCL25*) has been previously shown to play a critical role in the immune response to *Trichuris* (whipworm) by targeting lymphocytes to the small intestine ^37^. Infections by both intracellular and extracellular pathogens can also cause tissue damage; either from the antimicrobial Type 1 T helper (T_H_1) response which results in collateral tissue damage, or by physical trauma caused by helminths entering, exiting, and migrating through the host. As a result, wound healing and angiogenesis are critical components of the host response to infection ^38^. In addition to the increased wound healing mediated by over-expression of *IL5* ^39^, several other long-tailed goral PSGs are involved in these pathways and processes. Among them is the *colony-stimulating factor 2* (*CSF2*), which promotes the maturation of macrophages in lung and gut epithelium and is essential for the recruitment of innate immune cells to damaged tissues ^40^. We also observed seven other long-tailed goral PSGs in additional pathways involved in wound healing, including: Notch (*ITCRA*), WNT (*CSNK2B, FOSL1*, and *RUVBL1*), and NF-κB signaling (*BCL10, CSNK2B*, and *TNFAIP3*). Finally, we identified numerous PSGs involved in the T_H_1 response to viral and bacterial infections (Additional file 1: Table S6-S9). In particular, interleukin 12 (IL-12) is a cytokine that induces T_H_1 differentiation, and the ‘positive regulation of interleukin-12 production’ GO biological process (GO:0032735) was identified as statistically enriched (p-value: 0.01). Furthermore, the bactericidal/permeability increasing fold containing family A member 1 (BPIFA1) was positively selected, which is known to be expressed in the upper airways and plays a critical role in the inhibition of biofilm formation by pathogenic gram-negative bacteria such as *Pseudomonas aeruginosa* and *Klebsiella pneumoniae* ^41^.

To investigate adaptive evolution in the caprine common ancestor, we also analyzed the amino acid substitutions that were exclusively present in all members of the Caprini tribe (goat, sheep, mouflon, Tibetan antelope, long-tailed goral). From these, a total of 235 and 154 PSGs were detected using branch and branch-site models, respectively (Additional file 1: Table S9 and S10). Similar to the long-tailed goral, GO enrichment analyses of the Caprini PSGs showed numerous biological processes, molecular functions, and KEGG pathways related to both adaptive and innate immune function (Additional file 1: Table S11-S14). Of the PSGs in the innate response, the cytokines *interleukin 4* (*IL-4*) and *interleukin 33* (*IL-33*) are essential components of the T_H_2 immune response to helminths ^42-48^. Additional positively selected genes consist of other cytokines and chemokines, including *TNF, CXCL13*, and *CD34*, which play critical roles in the recruitment of immune cells to injured tissues ^49^. Furthermore, IL-17F recruits dermal innate immune cells during the inflammation and cell proliferation phases of wound response ^50^; and fibroblast growth factor1 (FGF1) is known to stimulate angiogenesis and tissue repair and to induce the proliferation of keratinocytes ^51^, which are required for the final stage of wound healing.

### Non-synonymous amino acid substitutions possibly confer selective advantages

The codon models for detecting positive selection may fail to detect genes that are positively selected if there are too few amino acid substitutions across a given gene ^52^. To overcome this limitation, we examined function-altering amino acid changes in single-copy orthologues to provide a more robust examination of the environmental adaptations of the long-tailed goral and Caprini. By aligning the 5,279 single copy orthologous genes from the eleven mammalian genomes, we investigated the amino acid substitutions among these species. We identified 3,802 unique amino acid substitutions (uAAS) in the long-tailed goral from 2,061 proteins (Additional file 1: Table S15). We then estimated the functional effects of these unique long-tailed goral substitutions using the programs PolyPhen2 ^53^ and PROVEAN ^54^. In total, PolyPhen2 predicted 1,582 (41.61%) amino acids as “possibly or probably damaging” and PROVEAN predicted 1,583 (41.63%) amino acids as “deleterious”. Of these, 1,341 (65.07%) proteins contain function-altering uAAS specific to the long-tailed goral as predicted by at least one program, and 967 long-tailed goral uAAS in 701 genes were predicted as function-altering substitutions by both programs. Similarly, we identified 6,543 uAAS unique to the Caprini lineage in a total of 2,744 proteins (Additional file 1: Table S16). Among them, 1,859 (28.41%) and 2,131 (32.08%) uAAS were predicted as function-altering by the PolyPhen2 or PROVEAN, respectively, in a total of 1,594 (58.09%) proteins with 1,004 uAAS in 674 proteins predicted by both programs.

To investigate the biological significance of these substitutions, we identified the KEGG pathways associated with the function-altering uAAS. In total, 187 (14.04%) proteins with unique long-tailed goral substitutions and 447 (51.68%) proteins with Caprini specific uAAS were mapped to 103 and 275 KEGG pathways, respectively (Additional file 1: Table S17). KEGG enrichment analyses of these two gene sets revealed a number of pathways related to immune function, RNA transport and biosynthesis, DNA repair, and cellular metabolism (Additional file 1: Table S18 and S19). Of particular interest in the Caprini are ‘Cytokine-cytokine receptor interaction’, ‘NF-κB signaling pathway’, ‘Apoptosis’, ‘Sphingolipid signaling pathway’, and ‘Complement and coagulation cascades’ (Table 1, Additional file 2: Figure S1). The pathway involved in ‘Cytokine-cytokine receptor interaction’ included 38 proteins containing unique Caprini uAAS. Of these, six additional tumor necrosis factor (TNF) family members were included over the three identified by the PSG KEGG enrichment (TNF, TNFRSF8, TNFRSF21). They were comprised of four TNF ligand superfamily members (TNFLSF4, 9, 10, and 15) and three TNF receptors superfamily members (TNFRSF1A, 9, 19). In addition, we identified five cytokines involved in hematopoiesis: the ciliary neurotrophic factor (CNTF), granulocyte colony-stimulating factor 3 (CSF3), interleukin 9 (IL9), and two interleukin receptors (IL23R, IL15RA); five chemokines: four C-C motif ligands (CCL4, CCL27, CCR1, CCR9) and one C-X-C motif ligand (CXCL9); and other minor cytokines: interferon kappa (IFNK), interleukin 26 (IL26), and interleukin 1 receptor type 1 (IL1R1). Also included as an enriched KEGG pathway related to immune function was ‘Complement and coagulation cascades’ (Additional file 1: Table S19). Our KEGG enrichment analyses of Caprini PSGs previously identified this pathway as enriched with four genes (*KNG1, MBL2, TFPI, CFI*) (Additional file 1: Table S13). Added to this, our functional analysis yielded an additional four proteins for complement cascade (C1QA, C1QC, C5AR1, and C8B) and seven proteins related to blood coagulation (SERPINC1, F5, F2RL2, PLAUR, PLAU, PLAT, and FGG), which may be an adaptation to steep and rugged habitats.

**Table 1.**
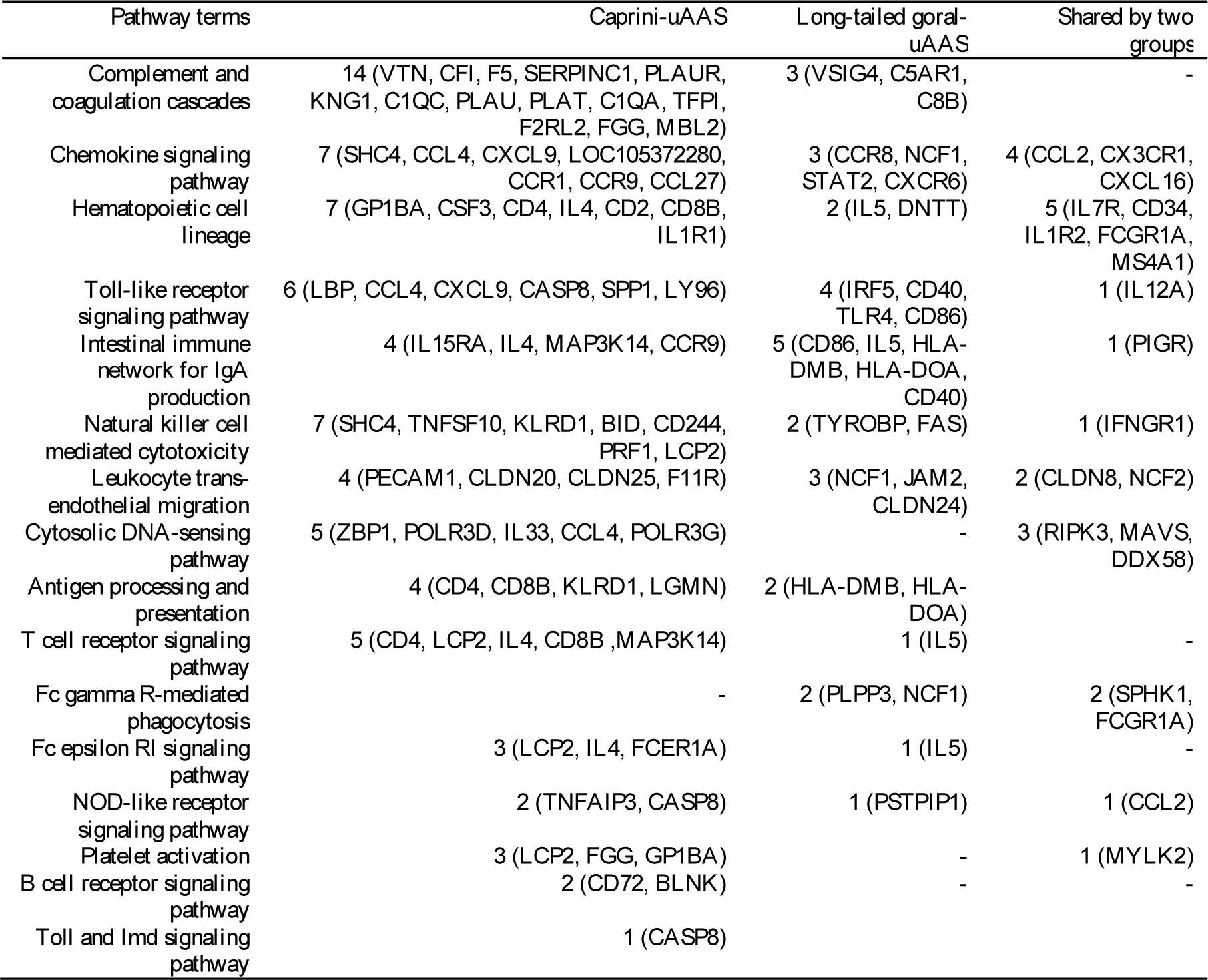
Immune response genes with function-altering uAASs

### Structural analysis of myostatin protein provides evidence for increased muscle development in Caprini

We observed nine Caprini-specific uAAS in MSTN, encoding the growth differentiation factor myostatin (Table 2). This protein is known to negatively regulate skeletal muscle mass by binding to the activin receptor type IIB (ACVR2B) ^55,56^, and humans and other mammals with loss of function mutations that block the activity of myostatin have significantly more muscle mass than those with functional proteins ^57-63^. To assess the impact of the Caprini uAAS on the function of this protein, we constructed a structural model of myostatin interacting with the receptor ACVR2B (Fig 3) and estimated the cumulative stability changes of the Caprini myostatin uAAS by calculating the change in binding free energy (ΔΔGBind) (Table 2). Among nine function-altering uAAS identified in Caprini, three amino acids physically interact with ACVR2B. Our simulations showed that the total binding energy was slightly increased by substituting these three Caprini uAAS. The resulting decrease in binding affinity between myostatin and ACVR2B in the Caprini may contribute to the increased muscle mass that has allowed this group to successfully inhabit steep, mountainous terrains; and warrants future experimental testing.

**Table 2.** Caprini-specific myostatin amino acids

| Position of<br>human<sup>a</sup> | Other<br>animals<sup>b</sup> | Caprini | $\Delta\Delta\text{GBind}^c$<br>(kcal/mol) | Interface | PolyPhen2<sup>d</sup> | PROVEAN<sup>d</sup> |
| --- | --- | --- | --- | --- | --- | --- |
| 16 | I | L | N/A |  | benign | neutral |
| 48 | T | N | N/A |  | benign | neutral |
| 113 | A | V | 0.25 |  | benign | deleterious |
| 130 | Q | E | 1.17 | O | benign | neutral |
| 133 | G | E | 0.56 |  | benign | deleterious |
| 148 | Y | H | 0.92 | O | benign | neutral |
| 247 | D | E | 0.64 | O | benign | neutral |
| 333 | R | K | 0.04 |  | benign | neutral |
| 366 | A | G | -0.34 |  | benign | neutral |

2 The position of the human myostatin protein sequence corresponding to caprini-specific amino acids<sup>b</sup> Other animals 3 represent non-Caprini animals analyzed in this study, including yak, bison, cattle, pig, horse, human.
<sup>c</sup> The negative sign in $\Delta\Delta\text{GBind}$ corresponds to a mutation predicted to increase the binding affinity.
<sup>d</sup> Programs estimating the function-altering AASs

**Figure 3.**
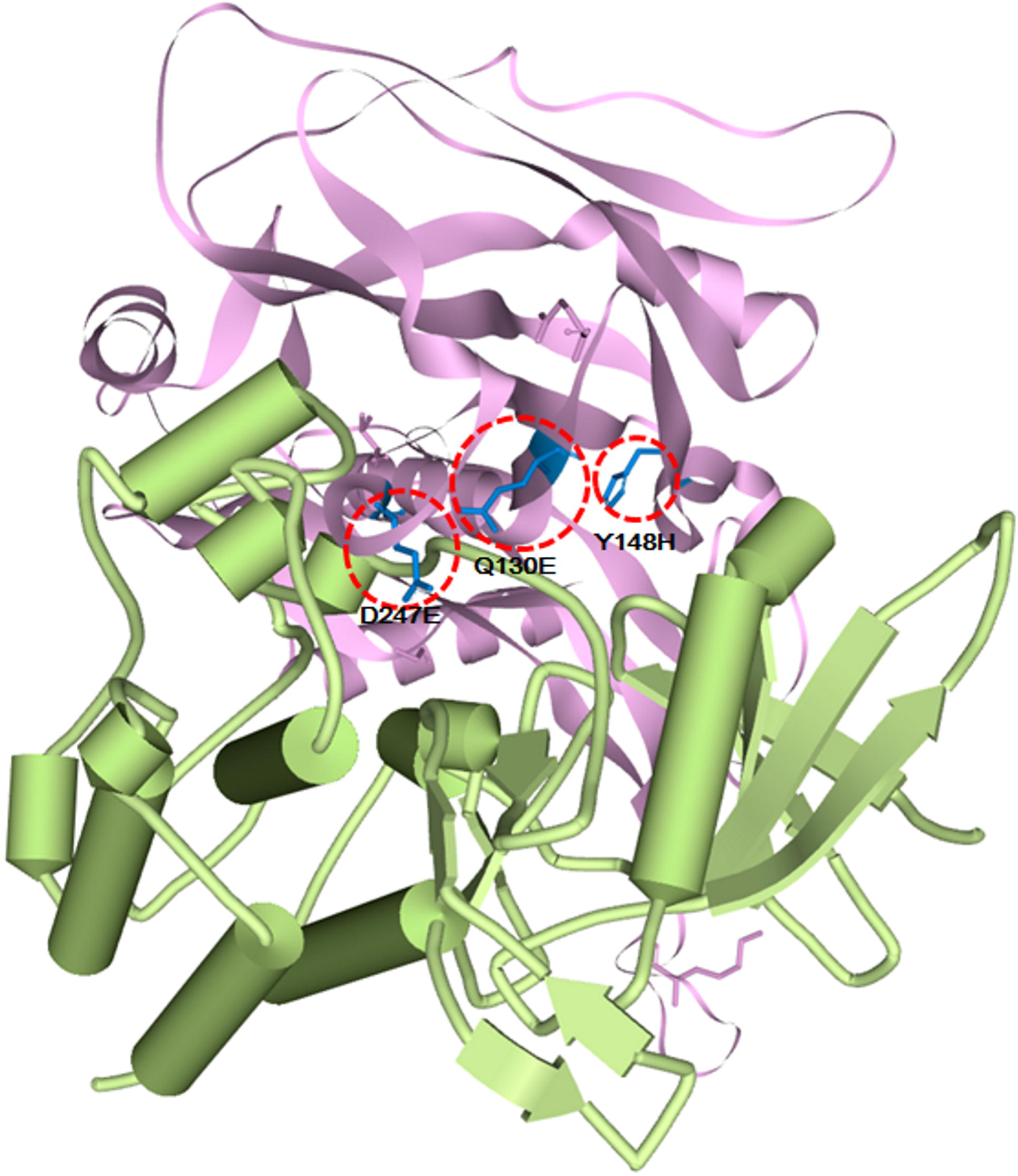
Caprini-specific amino acid change in the three-dimensional structure of ACVR2B and myostatin. The ACVR2B and myostatin are colored by green and purple, respectively. The five Caprini-specific amino acids that are located on non-interface region are shown in purple, and the three Caprini-specific amino acids that locate on interface region are in blue.

### Genetic diversity and population history

Deeply sequenced genomes allow for the identification of heterozygous positions and an estimation of the historical population structure ^64^. The nucleotide diversity for Caprini and Bovidae species was calculated by dividing the number of heterozygous sites by the genome size (Additional file 1: Table S20). The nucleotide diversity of the long-tailed goral was significantly lower (0.00113) than the mouflon (0.00379) and slightly higher than the formerly endangered Tibetan antelope (0.00079), which underwent a severe population bottleneck in the 1980’s ^65^. Interestingly, the nucleotide diversity of the long-tailed goral was also lower than the domestic sheep (0.00298) and goat (0.00189), which should be depressed from inbreeding associated with artificial selection (Additional file 1: Table S20). Effects, both environmental bottleneck and artificial selection, enable to decreased nucleotide diversities shown in Tibetan antelope and sheep and goat. A long-tailed goral, we applied in this study, lived in wild. Therefore, lower nucleotide diversity of the long-tailed goral was signature of inbreeding associated artificial selection. To further compare genetic diversity and infer the demographic histories of our five Caprini species, we used the Pairwise Sequentially Markovian Coalescent (PSMC) model ^66^. The population size of all of the Caprini species except for the long-tailed goral fluctuated over the past million years, while the goral showed an overall decline (Fig 4). Notably, the sheep and mouflon maintained a similar pattern until about 100,000 years ago, and the effective population sizes of these species both peaked approximately 1.0 MYA. The two domesticated Caprini (sheep and goat) showed secondary expansions at 100 KYA and 70 KYA, respectively, which were tightly linked with atmospheric surface air temperature (T_suf_); while the effective population size for the long-tailed goral and Tibetan antelope did not significantly increase around these times. In contrast, the effective population size of the long-tailed goral slightly increased about 30 KYA and the Tibetan antelope increased substantially, followed by declines in both species; though these date estimates occur at the edge of the accuracy range for this test. In conjunction with the observed population decline in this species, these findings suggest a possible need to review long-tailed goral’s IUCN conservation status and stress the importance of ongoing conservation and monitoring efforts.

**Figure 4.**
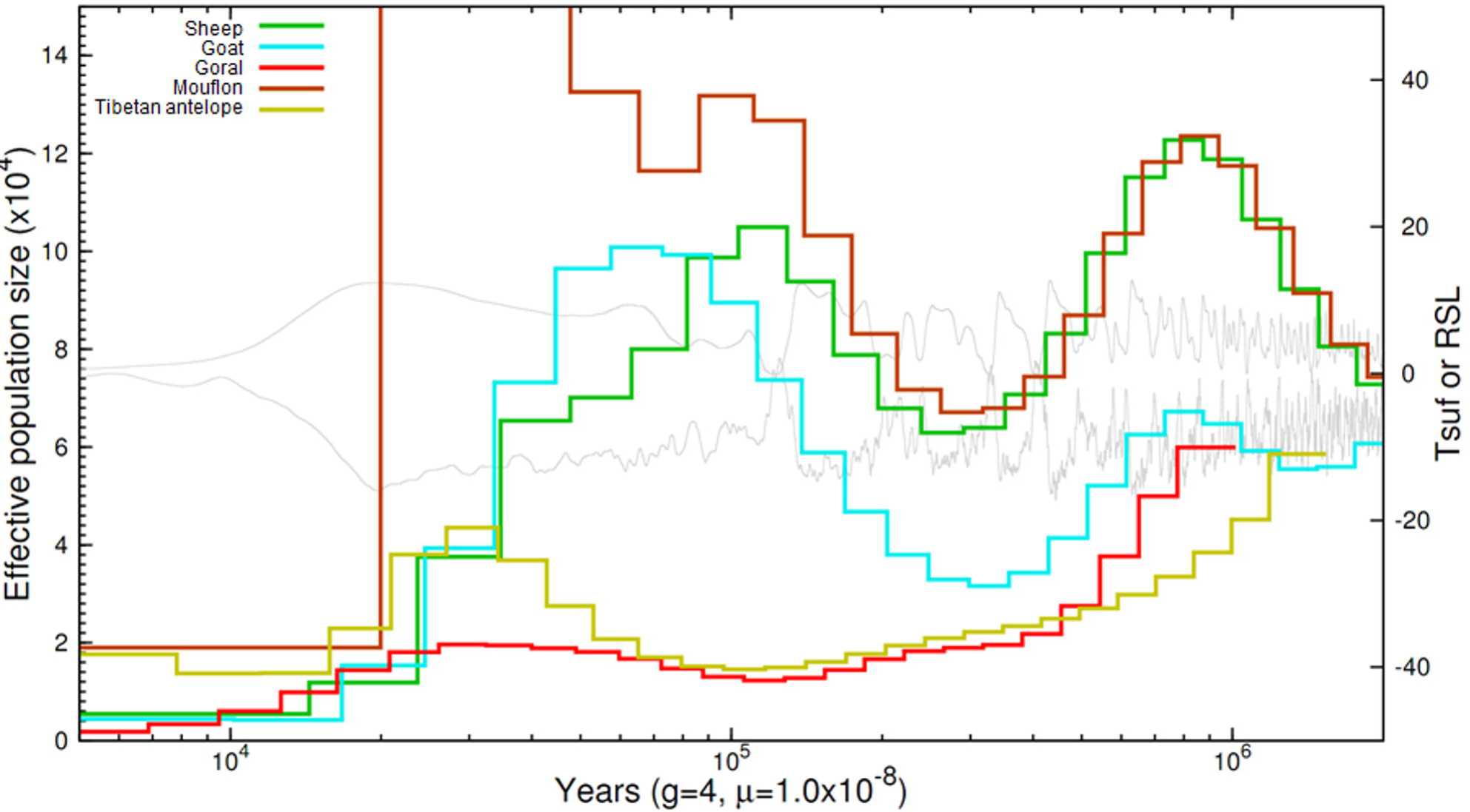
Demographic history of the Caprini. Demographic history of Caprini was analyzed by PSMC. g is generation time (years); μ, mutation rate per site per generation time; T_suf_, atmospheric surface air temperature; RSL, relative sea level; 10 m.s.l.e., 10 m sea level equivalent.

## Conclusions

We produced a whole genome sequence assembly of the endangered long-tailed goral by mapping 87 Gb of next-generation sequencing data to the closely related sheep genome. Using these data and ten publically available reference genomes, we produced a phylogeny dating the Antilopinae/Bovinae divergence to 14.1 MYA. Consistent with previous observations of serious infectious diseases and high parasite loads in long-tailed goral populations, evolutionary analyses of the long-tailed goral genome and the Caprini lineage identified positively selected genes in enriched GO and KEGG pathways related to immune function and blood coagulation; highlighting the importance of ongoing evolution in both the adaptive and innate immune responses in these groups. Together with previous mortality reports, these findings emphasize the need for additional studies to characterize the role of infectious disease and parasitism across extant populations of *N. caudatus*; which can be used to better inform management and conservation efforts. We hope that genome will stimulate comparative genomic studies aimed at investigating host-parasite co-evolutionary immune responses, potentially leading to targeted treatment strategies. We also identified the muscle development gene *MSTN* to be positively selected in the Caprini lineage, and discovered three amino acid substitutions that decrease the binding affinity between myostatin and its receptor ACVR2B. We speculate that these substitutions may have played a role in increasing muscle mass and, by proxy, climbing ability of the caprine common ancestor and warrants further functional investigation. Finally, we discovered lower levels of heterozygosity in the long-tailed goral than in other Caprini species, as well as a historical trend of population decline. These findings reinforce concerns over habitat fragmentation and genetic isolation in this species, and suggest that the IUCN status of the long-tailed goral should be re-evaluated. This foundational genome should stimulate future conservation genetic studies of this enigmatic endangered species.

## Methods

### Whole genome sequencing and alignment

A blood sample from a long-tailed goral was acquired from the Association of Korean Goral Conservation under the Cultural Heritage Administration 217 (Korea) Permit. Genomic DNA was extracted using a QiAamp DNA Mini kit (Qiagen, CA, USA) and quantified using an Infinite F200 Pro NanoQuant (TECAN, Männedorf, Germany). Libraries were prepared using an Illumina TruSeq kit (Illumina, CA, USA) according to the manufacturer’s instructions with an average insert size of 388 bp. 100 bp paired-end sequencing was completed using an Illumina HiSeq2000 with v. TrueSeq Kit v3 chemistry at the Korea Research Institute of Bioscience and Biotechnology (KRIBB). The raw sequence data were deposited in the Sequence Read Archive (SRP098771). The reads were aligned to the sheep genome assembly Oar_v4.0 using BWA-MEM (ver. 0.7.10) with default options ^67^. The rmdup command of SAMtools (ver. 0.1.19) was used to remove PCR duplicates ^68^, the GATK (ver. 3.3) IndelRealigner algorithm was used to realign reads ^69^, and SNVs were called using the mpileup command of the SAMtools with the default option followed by the view command with the -cg option. Finally, consensus sequences were generated using vcf2fq with -d 5 options in the SAMtools package.

### Orthologous gene clustering and phylogenomic analysis

Reference genomes for the goat (*Capra hircus*, CHIR_1.0), sheep (*Ovis aries*, Oar_v4.0), mouflon (*Ovis aries musimon*, Oori1), Tibetan antelope (*Pantholops hodgsonii*, PHO1.0), bison (*Bison bison bison*, Bison_UMD1.0), cattle (*Bos taurus*, Bos_taurus_UMD_3.1.1), yak (*Bos mutus*, BosGru_v2.0), horse (*Equus caballus*, EquCab2.0), pig (*Sus scrofa*, Sscrofa10.2), and human (*Homo sapiens*, GRCh38.p7) were downloaded from NCBI. To compare the long-tailed goral genes with protein sequences from these reference genomes, we constructed orthologous gene clusters using OrthoMCL 2.0.9 ^70^. To construct a phylogenetic tree, we used the 656,303 fourfold degenerate sites in the single-copy orthologous genes. A maximum likelihood phylogeny was then generated using RAxML ^71^ with the GTRGAMMA evolutionary model and 100 bootstrap replicates. And then, the divergence time was calculated using MEGA7 program ^72^ with maximum likelihood statistics and a Jukes-Cantor evolutionary model, along with the previously determined phylogenetic tree topology. The calibration times of horse-pig divergence (78.4 MYA) and human-horse divergence (97.5 MYA) were taken from the TimeTree database (http://www.timetree.org) ^73^.

### Identification of positively selected genes (PSGs)

The PRANK program was used to align the orthologous genes ^74^. The codon-based alignment was performed with this program, and the region containing the heterozygous SNVs was replaced with ‘NNN’. And the branch and branch-site models were used to identify PSGs. The CODEML program from the PAML 4.5 package was used to estimate the rates of synonymous (*d*_*S*_) and nonsynonymous substitutions (*d*_*N*_), and the general selective pressures acting on all species were estimated by applying the *d*_*N*_/*d*_*S*_ ratio (ω) for the branch model ^75^. PSGs were detected using the branch-site model using the *d*_*N*_/*d*_*S*_ ratio along each branch and applying a conservative 10% false discovery rate (FDR) for the likelihood ratio tests (LRTs) ^76^. To analyze the long-tailed goral PSGs, we set the genes of long-tailed goral as the foreground branch and those genes in other species were set as background. The foreground branch for the caprines included all five of the Caprini species (long-tailed goral, goat, sheep, mouflon, Tibetan antelope), with the six remaining species set as background. The DAVID bioinformatics resources tool was used to identify GO and KEGG pathway enrichment of the PSGs and uAAS ^77^.

### Functional effects of amino acid changes

uAAS were defined as amino acid substitutions that occurred in the target species, but no other species, and no gaps were allowed at the sites. Functional effects of the unique amino acid substitutions were predicted using PolyPhen2 ^53^ and PROVEAN v1.1 ^54^ programs with the default cutoff values, and the specific amino acids of the target species were substituted in the corresponding position of the human protein. KEGG pathways analyses were carried out using KAAS (KEGG Automatic Annotation Server) ^78^. As an additional functional analysis, we performed computer modeling of the myostatin protein structure using SWISS-MODEL ^79^, and a 3D structure of the ACVR2B protein was obtained from the PDB database (PDB ID: 2QLU) ^80^. The ZDOCK program ^81^ was used to construct the ACVR2B and myostatin complex protein structures, and the BeAtMuSiC program ^82^ was used to evaluate the change in binding affinity between two proteins.

### Demographic history analysis

To infer the population history of the long-tailed goral, we applied the Pairwise Sequentially Markovian Coalescent (PSMC) model for scaffolds ≥ 50 Kb in length. We performed 100 rounds of bootstrapping and used 1.0e-08 substitutions per site per generation as a mutation rate, and 4 years as a generation time as previously reported ^66^. Nucleotide diversity in the Caprini was calculated by dividing the number of heterozygous SNVs by the total sequence length of reference genome.

### Availability of supporting data

Whole-genome sequence data was deposited in the SRA database at NCBI with Biosample accession numbers SAMN06219541. The whole-genome sequence data can also be accessed through BioProject accession number PRJNA361026.

## Authors’ contributions

Conceived and designed the experiments: JB and WKP. Analyzed the data: OC, JK, and JAW. Study design, and sample preparation: DSK, JB, and WKP. Data interpretation: OC, JK, JAW, YSC, and JB. Wrote and edited the manuscript: OC, JAW, JK, JHJ, HL, SJ, YSC, SK, HL, SL, JSE, JAC, JA and JB.

## Additional files

**Additional file 1 – Table S1.** Sequencing and analysis statistics of the long-tailed goral’s WGS relative to the sheep genome. Table S2. Distribution of the homo-and hetero-SNVs for long-tailed goral in each sheep chromosome. **Table S3.** PSGs of the long-tailed goral using a branch model. **Table S4.** PSGs of the long-tailed goral using a branch-site model. **Table S5.** GO enrichment of long-tailed goral branch PSGs. **Table S6.** GO enrichment of long-tailed goral branch-site PSGs. **Table S7.** KEGG pathway enrichment of long-tailed goral branch PSGs. **Table S8.** KEGG pathway enrichment of long-tailed goral branch-site PSGs (Table S7). **Table S9.** PSGs of the Caprini using a branch model. **Table S10.** PSGs of the Caprini using a branch-site model. **Table S11.** GO enrichment of Caprini branch PSGs. **Table S12.** GO enrichment of Caprini branch-site PSGs. **Table S13.** KEGG pathway enrichment of Caprini branch PSGs. **Table S14.** KEGG pathway enrichment of Caprini branch-site PSGs. **Table S15.** Unique amino acid changes of the long-tailed goral. **Table S16.** Unique amino acid changes of the Caprini. **Table S17.** KEGG analysis of function-altering unique amino acid changes. **Table S18.** Enriched KEGG pathways using unique long-tailed goral amino acid substitutions. **Table S19.** Enriched KEGG pathways using unique Caprini amino acid substitutions. **Table S20.** Heterozygosity of the Caprini species.

**Additional file 2 - Figure S1.** Hotspots of the homozygous SNVs from the long-tailed goral distributed on the sheep genome. Hotspots were defined by the higher number of homozygous SNVs (pink for Z-score > 2 and red for Z-score > 3) in 0.5M window bins. The immune response genes with function-altering uAAS are given under the chromosomes; Caprini-uAAS (yellow), long-tailed goral-uAAS (green), and uAAS observed in both (blue).

