## Additional File 2 for "Genomic perspectives on adaptation and conservation in the endangered long-tailed goral (*Naemorhedus caudatus*)"

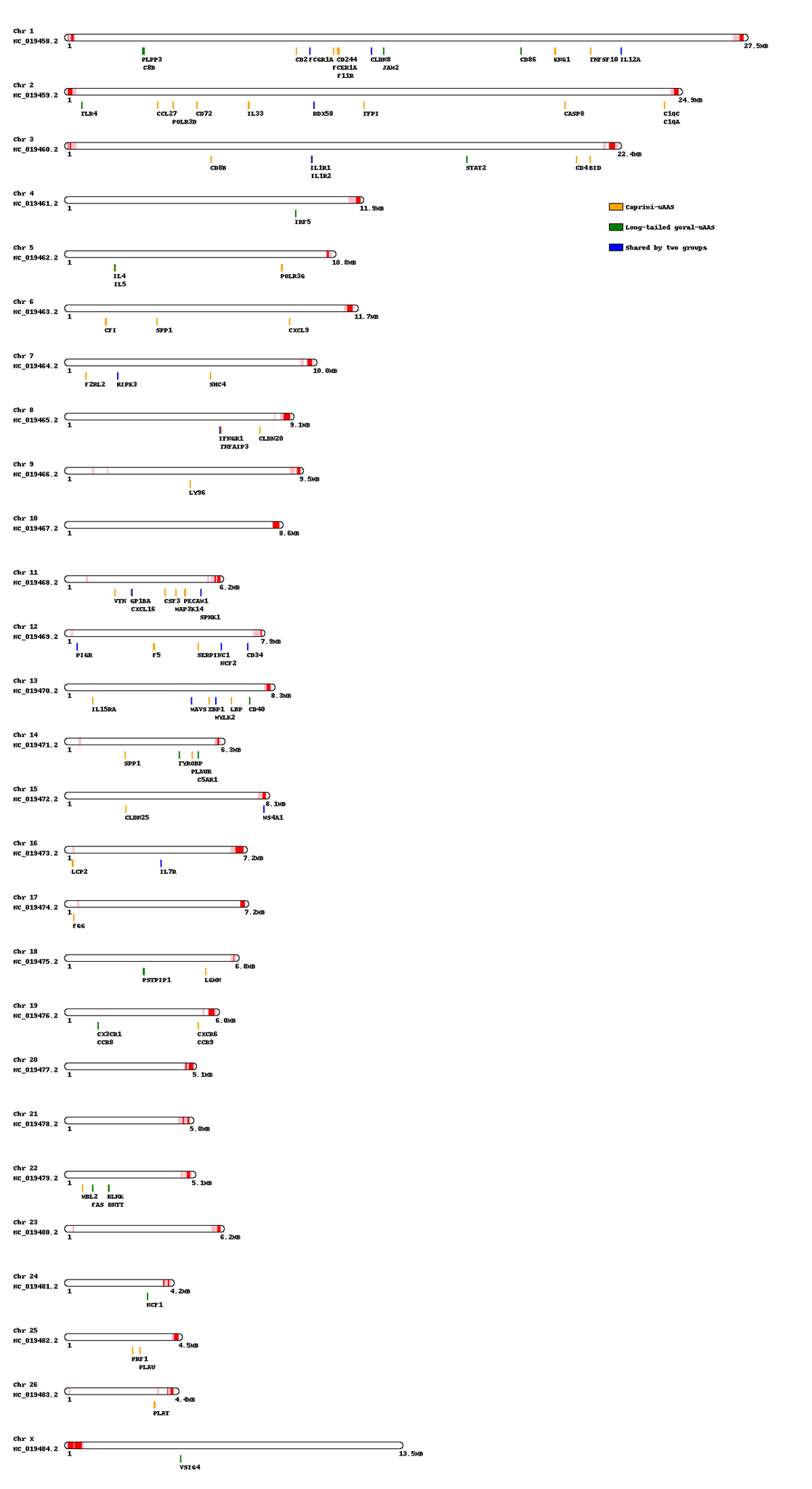

Figure S1. Hotspots of the homozygous SNVs from the long-tailed goral distributed on the sheep genome. Hotspots were defined by the higher number of homozygous SNVs (pink for Z-score > 2 and red for Z-score > 3) in 0.5M window bins. The immune response genes with function-altering uAAS are given under the chromosomes; Caprini-uAAS (yellow), long-tailed goral-uAAS (green), and uAAS observed in both (blue).
